# Fish functional diversity as an indicator of resilience to industrial fishing in Patagonia Argentina

**DOI:** 10.1101/2021.04.14.439740

**Authors:** Martha Patricia Rincón-Díaz, Nelson D. Bovcon, Pablo D. Cochia, María Eva Góngora, David E. Galván

## Abstract

The relationship between fish functional diversity and fishing levels at which its baselines shift is important to identify the consequences of fishing in ecosystem functioning. For the first time, we implemented a trait-based approach in the Argentine Patagonian sea to identify the vulnerability and spatiotemporal changes in functional diversity of fish assemblages bycatch by a trawling fleet targeting the Argentine red shrimp *Pleoticus muelleri* (Spence Bate, 1888) between 2003 and 2014. We coupled seven fish trophic traits to a reconstructed fish assemblage for the study area and bycatch and evaluated changes in fish species richness and four complementary functional diversity metrics [functional richness, redundancy, dispersion, and community trait values] along with fishing intensity, temporal use, latitudinal location, and depth of fishing grounds. Resident fishes larger than 30 cm in TL, with depressed and fusiform bodies, intermediate to high trophic levels, and feeding in shallow benthic, benthodemersal, and benthopelagic areas were vulnerable to bycatch. Fish assemblages exhibited a low functional trait redundancy, likely related to species influxes in a biogeographic ecotone with tropicalisation signs. Significantly increases in fish trait richness and dispersion polewards and with depth suggested new functional roles in these grounds, matching trends in community body size, reproductive load, maximum depth, trophic level, and diet breadth. Finally, a temporal increase in fish species and functional trait removal in fishing grounds led to trait homogenisation since the first year of trawling. The identified tipping points in temperate fish functional trait diversity highlight trait-based approaches within ecosystem-based fisheries management.

## Introduction

Ecosystem-based fisheries management claims, among other issues, a shift towards monitoring and identifying fishing practices that allow balanced exploitation of fisheries resources while protecting vulnerable components of ecosystem functioning (Zhou et al., 2010). While achieving this balance depends on a trade-off between fisheries targets and vulnerability of non-target species to fishing gears, trait-based approaches appear as an option to identify traits that place species prone to incidental captures and effects of trait removal in functional community structures (Barnett et al., 2019). Functional traits are morphological, behavioural, physiological, and life history attributes of organisms that determine their fitness through habitat filtering and help scientists describe species’ ecological roles in a system (Diaz and Cabido 2011; Mouillot et al., 2013a; Villéger et al., 2017). Traits correlate to ecosystem processes (e.g., food web controlling through predation) and their dependent services (e.g., food provision for humans) (Cadotte et al., 2011, 2017; Diaz and Cabido 2011; Hooper et al., 2005; Villéger et al., 2017), and also provide a common currency to evaluate marine communities resilience [recovery sensu] to habitat disturbances through species trait presence and abundance (Hodgson et al., 2015; Oliver et al., 2015; van der Linden et al., 2016). For example, some sharks, located at the top of food webs, may control marine communities’ structures and energy fluxes throughout fish and invertebrate consumption (Heaithus et al., 2008). Sharks grow slowly, mature late and have low fecundity, which prone them to be functionally extinct by overfishing with gillnets and long lines and inadequate fishing regulations in many global reefs (MacNeil et al., 2020). By linking species traits to disturbances, ecological studies can understand species loss’s consequences in functional diversity (Caddotte et al., 2017; Nyström, 2006).

Fishing acts as a selective driver, targeting and incidentally removing particular fish trait combinations depending on gears (Koutsidi et al., 2016) and influencing the functional structure of fish assemblages. While purse-seine gears remove small pelagic fishes that transfer energy from plankton to higher trophic levels, trawling fisheries pull fish traits associated with the seafloor, large body size, cryptic exposure, and ambushing behaviour (Koutsidi et al., 2016). Trawling also produces biomass reductions in sedentary, long-live, late mature, and low fecund fishes such as Chondrichthyes (Henriques et al., 2014), removes morphological traits related to low propulsion (Mouchet et al., 2019) and redundant species with intermediate to high trophic levels, benthopelagic habits, and fusiform body shapes (Herrera Valdivia et al., 2016). The trait composition of fish assemblages can respond to environmental conditions rather than fishing as well. In fact, in the north sea, beam trawling had a weak association with removing traits such as life history, growth rate, reproduction, and trophic levels, being the spatial fish trait community composition determined by depth (Beukhof et al., 2019). However, fishing selectivity is related to long-term body size changes (Beukhof et al., 2019). Because functional traits respond to environmental filtering and disturbances, trait-based approaches are essential tools in productive fishing areas, such as the temperate Southwest Atlantic Ocean, to map fish trait distributions and understand the fish trait-environmental relationships (Beukhof et al., 2019). Moreover, these approaches are helpful to understand fish vulnerability to different catch sources (Koutsidi et al., 2016; Mouchet et al., 2019; Nash et al., 2017).

Fisheries in the Argentine Patagonian Sea, southward -41° S, are potential drivers of change in fish community species composition and food web structure, but their effects on the fish functional trait diversity are unstudied yet. The Argentine hake *Merluccius hubbsi* Marini 1933, red shrimp *Pleoticus muelleri* (Spence Bate 1888), and squid *Illex argentinus* (Castellanos 1960) fisheries in Patagonia contribute to more than 75% of the current annual landings in Argentina, with exports raised 1520 million dollars deal in 2019 (MAGyP, 2020). Industrial fisheries of M. *hubbsi* and *P. muelleri* incidentally capture 101 fish species, including 69 Osteichthyes, 29 Chondrichthyes, and three Agnatha (Bovcon et al., 2011, 2013; Gongora et al., 2009, 2020; Ruibal, 2020), and fish species with partial retraction in their geographic distribution ranges coincide with commercial fishing targets (Galván et al., in press). The trophic structure of demersal fish assemblages in the San Jorge Gulf (SJG), one productive area in the region, also changed due to industrial fishing. Fish community diet breadth and trophic levels increased in the last decade due to the inclusion of fisheries discards in the intermediate predators’ diet (Funes et al., 2019; Funes, 2020). The elasmobranchs’ proportional biomass to teleosts and fish community body size decreased in the gulf due to the high bycatch of large chondrichthyans by industrial trawling (Funes, 2020). Despite the regional advances in understanding fishing effects on the taxonomy, trophic and size structure of fish communities, descriptions of functional traits for local fish assemblages exist. Neither identified spatiotemporal patterns in their functional diversity or fishing levels at which its baselines shift.

Here, we implemented a synthesis of existing information from local studies and onboard observers programs to identify fish trophic traits removed incidentally by industrial fishing and understand its effects on the fish ecological functions. To achieve these goals, we identified the fish vulnerability to incidental capture by the fishery of the shrimp *P. muelleri* based on differences of trait values by species occurrence and observation in the bycatch of the San Jorge Gulf (SJG) and adjacent waters in Argentina between 2003 and 2014. We also evaluated the spatio-temporal changes of four complementary metrics of functional fish diversity [functional richness, redundancy, dispersion, and weighted mean trait values] in bycatch along gradients of fishing intensity temporal use, latitudinal location, and depth of fishing grounds. Given that fisheries produce changes in the abundance and functional trait composition of fish assemblages, we worked under the hypothesis that there are differences in functional diversity metrics along the trawling intensity gradient and temporal use of fishing grounds. We expect descriptors of fish taxonomic and functional richness and dispersion in the bycatch to increase with high fishing intensity because it increases the probability of capturing species from fish assemblages. However, we expect a decline along with fishing grounds’ temporal use due to the low species selectivity of trawling activities that erode trait redundancy in fish assemblages. Finally, since latitude relates to marine fish species’ biogeographic distribution in the study area, we expect to observe an increase in fish taxonomic and functional richness and dispersion towards lower latitudes.

Research on fish communities’ ecological responses to disturbances is essential because it allows us to understand the footprint produced by fishing in marine ecosystems. The differences between fish functional diversity metrics along the fishing effort gradient will allow us to identify 1) fish functional traits removed by the fishery, 2) ecological responses of the fish assemblages to fishing, and 3) areas of high functional trait diversity that may require protection in the study area. Our characterisation of functional diversity in fish assemblages and the description of its responses to incidental fishing is the first one in Argentina. This information is essential to establish thresholds of sustainable exploitation of marine resources while maintaining the trophic structure of associated biotic communities to industrial fisheries. Our study also supports the application of trait-based approaches to ecosystem-based management in high productivity areas in the Southwest Atlantic Region.

## Materials and methods

### Study area and characterisation of incidental fishing

Our evaluation used a bycatch database for the fishery of the shrimp *P. muelleri* collected by the Fishery Observer Program from the Fisheries Secretariat of the Chubut Province, Argentina, on board the double-rigged otter trawler fleet. Data collection was conducted under the law XVII-70 of the Chubut Province to monitoring and control the use of marine resources (http://www.legischubut.gov.ar/hl/digesto/lxl/XVII-70.html). No fish was handled, sampled, or used in experiments during this study. The program registered the fleet’s bycatch of 38.676 hauls deployed between -41□ and -48□ S and -62□ and -67□ W in the San Jorge Gulf (SJG), within the Chubut and Santa Cruz Provinces’ jurisdiction, and adjacent waters of National jurisdiction in the Southwestern Atlantic Ocean between February and November from 2003 to 2014. The SJG, the largest gulf on the Argentine continental shelf, covers 39,430 km^2^ from -45□ to - 47□ S and -68□ to -66□ W (Matano and Palma, 2018). It reaches a maximum depth of 100 meters at its central basin and exhibits a shallower bank at its eastern limit with the open shelf (60 to 90 m depth) (Matano and Palma, 2018). Its high productivity supports the fisheries of *M. hubbsi* and *P. muelleri* due to the dominance of westerlies winds and strong seasonal ocean circulation with nutrient-rich waters of the Magellan Straits that generate a thermohaline front at the gulf’s southern end, developing a loop around it in winter (Glembocki et al., 2015; Palma and Matano, 2012). Tidal fronts are also identified in the gulf’s southern shelf with a peak in summer and at the northern side during spring and autumn due to the interaction between tides and vertical stratification of the water column (Glembocki et al., 2015). The studied area is located in the ecotone between the Argentine and Magellan Biogeographic Provinces, which influence fish distribution with species influxes from warm and cold-nutrient rich waters from subtropical and sub-Antarctic origins, respectively (Balech and Erlich, 2008; Figure 1a).

**Figure 1.**
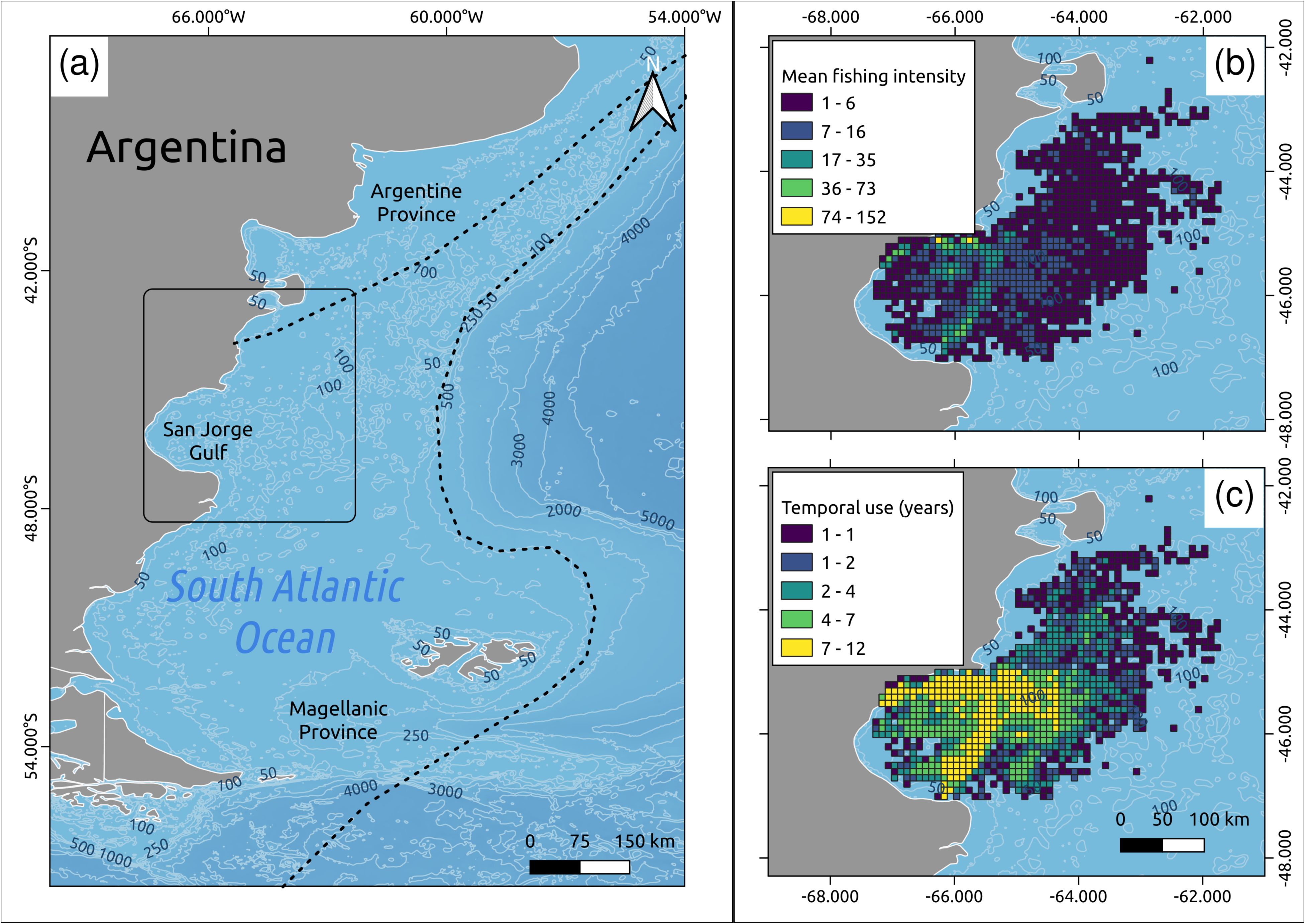
The double-rigged otter trawler fleet’s operational area in the San Jorge Gulf and adjacent national waters between 2003 and 2014 (rectangle in map A). Coloured maps denote the mean annual haul density (B) and total temporal use of fishing grounds by the fleet during the studied period (C).

The fleet ceased operations in the SJG in 2017 when the fishery of *P. muelleri* was closed in this area and then focused its activities in the adjacent national waters. A total of 90 vessels with 23 to 53 m in length comprise the fleet. Each vessel has a trawl net in each band of 45 mm mesh size on the bag and a vertical and horizontal opening of 1.5 m and 30 to 50 m, respectively (Gongora et al., 2012; Roux et al., 2007). Each observer onboard collected information on species incidentally captured at the haul level. They also registered the vessel type and hauled date, depth, time, and geographic location. Observers calculated the number of individuals per identified fish species in the total catch to assign a relative abundance category called observation percentage. Four categories were assigned: dominant (> 50% of individuals), abundant (25 to 50%), common (5 to 25%); and rare (< 5%) (Bovcon et al., 2013; Gongora et al., 2009).

### Fishing effort indicators

We used the mean annual haul intensity and years of use of fishing grounds to characterise the fleet’s spatial and temporal fishing effort (Figure 1b). The fishing area was divided into 6’ x 6’ cells (11 x 11 km^2^) to compare our results with previous fishing intensity descriptions for the otter trawler fleet in the area (Gongora et al., 2020; Ruibal Nuñez, 2020). The haul intensity was estimated considering only the initial location of each haul within a cell, and we think this is a reliable indicator of the spatial distribution of fishing grounds based on feasible data obtained from the fishing fleet. Haul intensity also described trawling intensity effects in spatial patterns of fish functional diversity in the Mediterranean sea (Henriques et al., 2014). Haul intensity ranged from one to 150 hauls per cell during the 12 evaluated years. Cells with high haul intensity were considered areas of considerable cumulative spatial fishing effort by the fleet. We characterised the temporal use as the number of years that the fleet used a cell at the haul evaluation (Figures 1c and S3). The minimum haul intensity considered was one haul per year, and the cell use ranged from one to 12 years without homogeneous spatial use within the area. Within the SJG, some fishing areas reached 12 years of use, and national waters had less than five years.

### Reconstruction of fish assemblage species composition

We reconstructed the fish assemblage for the entire study area through an extensive literature review on species occurrences in the Argentine continental shelf between the 42° and 48°S, and up to 100 m depths, including benthic and pelagic zones within the fleet’s operational area (Figure 1). Studies dated between 1978 to 2020 and included information on incidental fishing from the fisheries of *P. muelleri*, *M. hubbsi*, anchovy *Engraulis anchoita* Hubbs & Marini, 1935, independent surveys, and ichthyologic collections. The community reconstruction went through a review by local experts in fish biology to confirm species presence and assign an occurrence categorisation as a resident, seasonal, occasional, and anecdotal in the study area. A total of 127 marine fish species were recorded for the area (Table S2).

### Compilation of functional traits

A literature review of 327 regional studies was conducted to compile morphological, behavioural, and life-history traits that describe the trophic function of the 127 fish species recorded in the entire study area (Table 1, Figure S2, Appendix S2). The geographic scope of reviewed studies included our study area and the Argentine continental shelf primarily. We used trait values from Fishbase.org (Froese and Pauly, 2020) only when regional data were not available. All selected traits are related to the food web control and resources use, and resilience of fish communities to fishing (Beukhof et al., 2019; D’aganta et al., 2016; Koutsidi et al., 2016; Maureaud et al., 2019b; Michelli et al., 2004, 2014; Mouchet et al., 2019; Rincon-Diaz et al., 2018). Compiled traits included the location of fish species in the water column, body shape, and maximum regional depth that relate to species habitat use and strategies for feeding (Beukhof et al. 2019; Michelli et al. 2014; Mouillot et al. 2014: Rincon-Diaz et al. 2018; Stuart-Smith et al. 2013). We also included the fish trophic level and diet breadth that describe the species position in the trophic web and their omnivory capacity, respectively (Beukhof et al., 2019; D’agata et al., 2016; Mouillot et al. 2014, Rincon-Diaz et al. 2018; Stuart-Smith et al. 2013). We included regional maximum body size as a proxy of the amount of energy transported by a species in the food web and to describe fish species vulnerability to fishing or their facility to escape from fishing gears, which increases with small body sizes (Mellin et al. 201; Michelli et al. 2014; Mouillot et al. 2014; Mouchet et al. 2019; Stuart-Smith et al. 2013). The reproductive load represented a trade-off between species’ energetic investment between growth span and maturation, with a negative relationship with fish size (Tsikliras and Stergiou, 2013). The reproductive load also showed chondrichthyans’ vulnerability to trawling activities in our study area (Ruibal Nuñez, 2020). The trophic level was calculated with the TrophLab software (Pauly et al., 2000) using diet content analyses conducted primarily in the Central Patagonian Region of Argentina. The diet breadth described the number of taxonomic families or higher-order groups consumed by a fish species (Froese and Pauly, 2020). The reproductive load was calculated as the ratio between the average length at first maturity and the maximum length reached by a population (Tsikliras and Stergiou, 2013). We considered trait category combinations to account for intraspecific variability in functional traits (Rincon-Diaz et al., 2018). The water column position and body shape were categorical traits. The fish body size, reproductive load, maximum depth, trophic level, and diet amplitude were scaled by their maximum value to improve linearity in the ordination and were also categorised to calculate the number of functional entities and their redundancy.

**Table 1.**
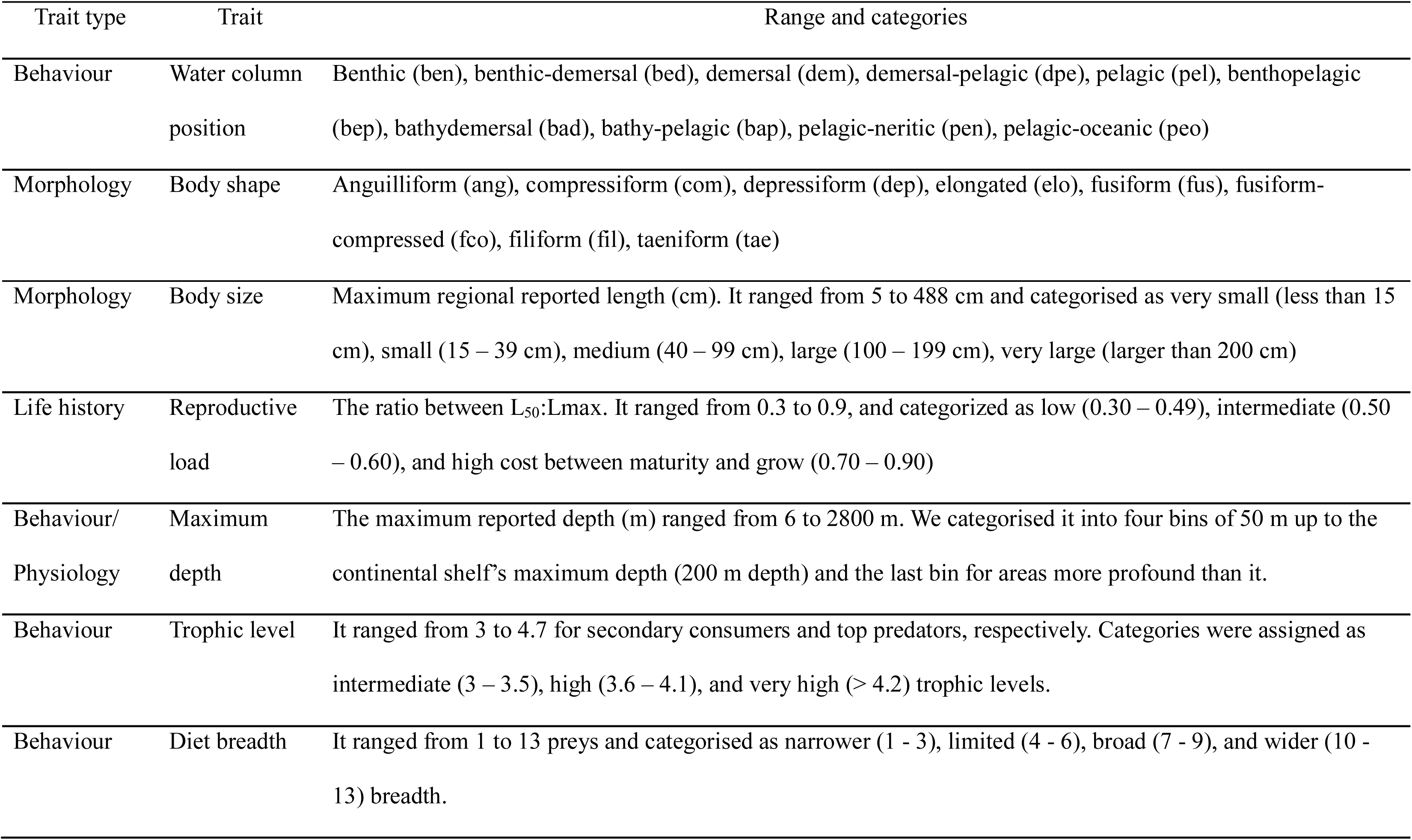
Functional traits used to describe the trophic function of marine fish species in the study area.

| Trait type | Trait | Range and categories |
| --- | --- | --- |
| Behaviour | Water column position | Benthic (ben), benthic-demersal (bed), demersal (dem), demersal-pelagic (dpe), pelagic (pel), benthopelagic (bep), bathydemersal (bad), bathy-pelagic (bap), pelagic-neritic (pen), pelagic-oceanic (peo) |
| Morphology | Body shape | Anguilliform (ang), compressiform (com), depressiform (dep), elongated (elo), fusiform (fus), fusiform-compressed (fco), filiform (fil), taeniform (tae) |
| Morphology | Body size | Maximum regional reported length (cm). It ranged from 5 to 488 cm and categorised as very small (less than 15 cm), small (15 – 39 cm), medium (40 – 99 cm), large (100 – 199 cm), very large (larger than 200 cm) |
| Life history | Reproductive load | The ratio between $L_{50}$:$L_{max}$. It ranged from 0.3 to 0.9, and categorized as low (0.30 – 0.49), intermediate (0.50 – 0.60), and high cost between maturity and grow (0.70 – 0.90) |
| Behaviour/<br>Physiology | Maximum depth | The maximum reported depth (m) ranged from 6 to 2800 m. We categorised it into four bins of 50 m up to the continental shelf's maximum depth (200 m depth) and the last bin for areas more profound than it. |
| Behaviour | Trophic level | It ranged from 3 to 4.7 for secondary consumers and top predators, respectively. Categories were assigned as intermediate (3 – 3.5), high (3.6 – 4.1), and very high (> 4.2) trophic levels. |
| Behaviour | Diet breadth | It ranged from 1 to 13 preys and categorised as narrower (1 - 3), limited (4 - 6), broad (7 - 9), and wider (10 - 13) breadth. |

### Fish functional vulnerability to the bycatch

We evaluated the categorical traits with a Chi-square analysis and numerical traits with a paired non-parametric Mann–Whitney U test to identify if the fish functional trait composition in the entire study area responded randomly to the species occurrence status, inclusion in the incidental capture, and percentage of observation within the bycatch. Tests were run with the chisq.test and Wilcox.test functions in the R software (R Core Team 2020). We used Bonferroni significant corrections with statistical significance at p-value < 0.05 in the Mann-Whitney U tests.

### Calculation of fish functional diversity and composition

We calculated taxonomic species richness [SR], and functional richness [FRic], redundancy [FRed], and dispersion [FDis] to complementary describe the diversity and redundancy of trophic roles in fish assemblages of the entire studied area and bycatch. Functional diversity values for the whole studied area were reported as “reference values” in a non-fishing scenario. We selected these three complementary descriptors of functional diversity because they describe the functional organisation of communities in terms of diversity [FRic], variation [FDis], and redundancy [FRed] of ecological roles to accomplish an ecosystem function (Mouillot et al., 2013) and significantly complement the information provided by other functional diversity metrics (Appendix S1, Table S3). The FRic measures how much of the niche space a community occupies by estimating the convex polygon volume formed by species in the trait space (Mouillot et al., 2013). High FRic values indicate many functional roles in the community (Villéger et al., 2008). Functional redundancy describes the mean number of species that accomplish a particular role by considering the trait combinations within a community (Mouillot et al., 2014). Values higher than one suggest that more than one species performs a functional role, increasing the community resilience against species loss (Mouillot et al., 2014). Functional dispersion describes the variability of functional roles in a community. It provides an understanding of trait redundancy by weighting species relative abundances and their pairs dissimilarity distances to the centroid of the trait space (Laliberté and Legendre, 2010; Laliberté et al., 2014; Schleuter et al., 2010; Villéger et al., 2008). High values of functional dispersion suggest a high variability in functional roles, and low values indicate that more than one species provides a specific ecosystem function, increasing the community resilience to disturbances (Mouillot et al., 2013; Rincón-Díaz et al., 2018).

Previous to calculating metrics, we examined the relationship between the annual sampling effort by the onboard observers’ program and cumulative species richness and identified that the bycatch’s fish community was satisfactorily characterised (Figure S1). We followed Villéger 2020 to calculate functional diversity metrics (http://villeger.sebastien.free.fr/Rscripts.html). We used the compiled trait information to calculate a pairwise Gower distance matrix between the fish species recorded in the entire area and a Principal Coordinate Analysis (PCoA) based on the distance matrix to obtain species coordinates in the trait space (Mouillot et al., 2014; Villéger et al., 2008, 2017). We selected the first four PCoA axes to calculate functional diversity metrics because they closely represented the species pairwise Gower distances (D’agata et al., 2016; Marie et al., 2015; mean squared deviation, mSD = 0.0018). We used species coordinates to calculate species richness and functional richness and redundancy for fish assemblages in the entire study area and each haul. We also used the coordinates to calculate functional dispersion in each haul by using an adapted value within each original interval for the observation percentage of species in the bycatch following an increasing scale. Original intervals of observation were rare (< 5%), common (5 - 25%), abundant (25 - 50%), and dominant (> 50%) (Góngora et al., 2020). We classified final species observation in the bycatch as rare (2.5%), common (12.5%), abundant (25%), and dominant (50%). We calculated the Gower distance matrix, PCoA, and functional diversity descriptors using the species_to_FE, FE_metrics, quality_funct_space, plot_funct_space, multidimFD functions by Villéger (2020) in the R software (R Core Team 2020).

We also calculated community weighted mean trait values (CWM) at the haul level to explore spatiotemporal changes in the fish trait community composition in the bycatch. CWM values provide information about the dominant trait values in a community by considering relative species abundance (Laliberté et al., 2015). We calculate CWM by using the *functcomp* function from the FD package in the R software (Laliberté et al., 2015).

### Spatiotemporal patterns of fish trait diversity and composition

We evaluated the relationship between fish functional trait diversity and composition in the bycatch of the red Argentine shrimp industrial fishery with Generalised Linear Mixed Models (GLMM). Response variables were the fish species richness, functional richness, redundancy, and dispersion, and continuous community weighted mean trait values in the bycatch. Predictors of the otter trawler fleet were the mean fishing intensity, temporal use of fishing grounds, and latitude and depth of hauls deployed. We included surveyed years and observers on board the fleet as random effects accounting for their temporal variation. We followed the protocol proposed by Zuur et al. (2010) for data exploration and Zuur et al. (2009, 2016) for modelling and results presentation. We kept all predictors because they did not show high significant collinearity (Rho < 0.60 or > -0.60, and p-values > 0.05, Table S4) and removed outliers of response variables to improve linearity in models. We ran seven models for each response variable with single and combined predictors. We modelled the taxonomic species richness with a Poisson distribution, functional richness and dispersion, fish community body size and diet breadth with a Gaussian distribution, and trophic level with a Gamma distribution using the lme4 package in the R software (Bates et al., 2015). Functional redundancy was not modelled because it showed a regular distribution and no variability in data. We based model selection on predictors with variance inflation factors (VIFs) < 2, the lowest Akaike’s information criterion (AIC), and delta AIC < 2 (Zuur et al., 2009). We used the car package to calculate the VIFs in the R Program (Fox, 2019). Mean estimates of fixed predictors were reported for the best model. We visually inspected the differences between functional diversity metrics along the trawling intensity gradient with reference values for the entire studied area to understand how close the fleet is to the functional capacity of fish assemblages in the system. Changes in dominant categorical traits along predictors were analysed with non-parametric Mann-Whitney pairwise tests with Bonferroni corrected p-values (< 0.05) by using the *pairwise.wilcox.test* function from the stats package in the R software (R Core Team 2020).

## Results

### Fish functional trait composition and vulnerability to the bycatch

We found that fish trait composition was related to species occurrence in the entire study area and species inclusion and abundance in the bycatch. Fish water column position and body shape varied by these three species status (Figure 2, Chi-square statistic < critical value, df_position_ = 9, df_shape_ = 8, p-value < 0.05). At least 50% of the area’s resident fish species were benthic and benthodemersal with depressed, elongated, and fusiform body shapes. Seasonal species were mainly benthic and benthopelagic with fusiform bodies; occasional species were mainly benthopelagic with compressed and fusiform bodies; and anecdotal species were pelagic and pelagic-oceanic with fusiform bodies (Figure 2a,d).

**Figure 2.**
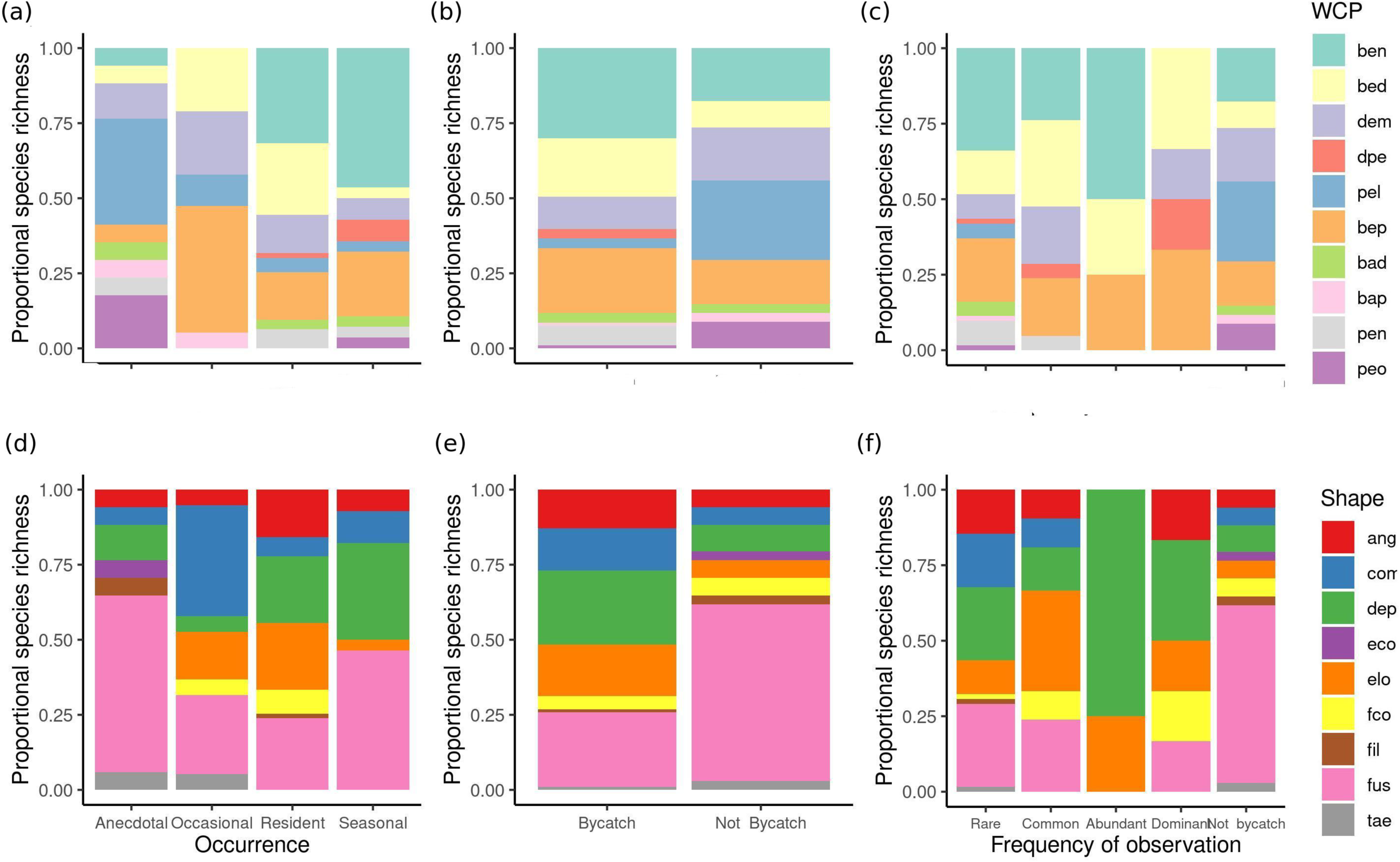
Proportional species richness by categories of fish water column position (WCP) and body shape. Trait composition was described by fish species occurrence (a,d), and inclusion (b,e), and percentage of observation in the bycatch of the otter trawler fleet (c,f).

By considering species included in the bycatch, more than 50% of caught fishes were benthic and benthopelagic with depressed, fusiform, and compressed bodies, and those not caught were mainly fusiform fishes with pelagic, benthic, and demersal feeding habits (Figure 2b,e). More than 50% of the bycatch’s dominant species were benthodemersal and benthopelagic fishes with depressed, elongated, and fusiform body shapes (Figure 2c,f). The majority of abundant species were benthic fishes with depressed bodies; common fishes were benthic and benthodemersal with elongated and fusiform bodies, and rare species were benthic and benthopelagic fishes with fusiform and depressed bodies.

We found trait value differences among species only for the maximum reported depth based on fish species occurrence in the study area and for body size and trophic level by the combination of species occurrence and observation percentage in the bycatch (Figure 3a-d, paired Mann– Whitney U test, p-value < 0.05). Fish species not caught by the fleet were residents with smaller body sizes (< 32 cm TL), trophic levels (< 3.5), and maximum depths (< 55 m) than other residents in the bycatch and corresponded to the family Atherinopsidae (the silversides *Odontesthes incisa* (Jenyns 1841) and *O. nigricans* (Richardson 1848)), Clinidae (*Ribeiroclinus eigenmanni* (Jordan 1888)), Nototheniidae (the Patagonian rock cod *Patagonotothen brevicauda* (Lönnberg 1905), *P. cornucola* (Richardson 1844), and *P. sima* (Richardson 1845)), Tripterygiidae (the Cunningham’s triplefin *Helcogrammoides cunninghami* (Smitt 1898)), and Zoarcidae (*Crossostomus fasciatus* (Lönnberg 1905)). In this context, we identified vulnerable species to the otter trawler fleet as resident species with body sizes larger than 30 cm TL, intermediate to higher trophic levels, and associated to benthic and benthopelagic areas between 50 to 500 m depth for feeding.

**Figure 3.**
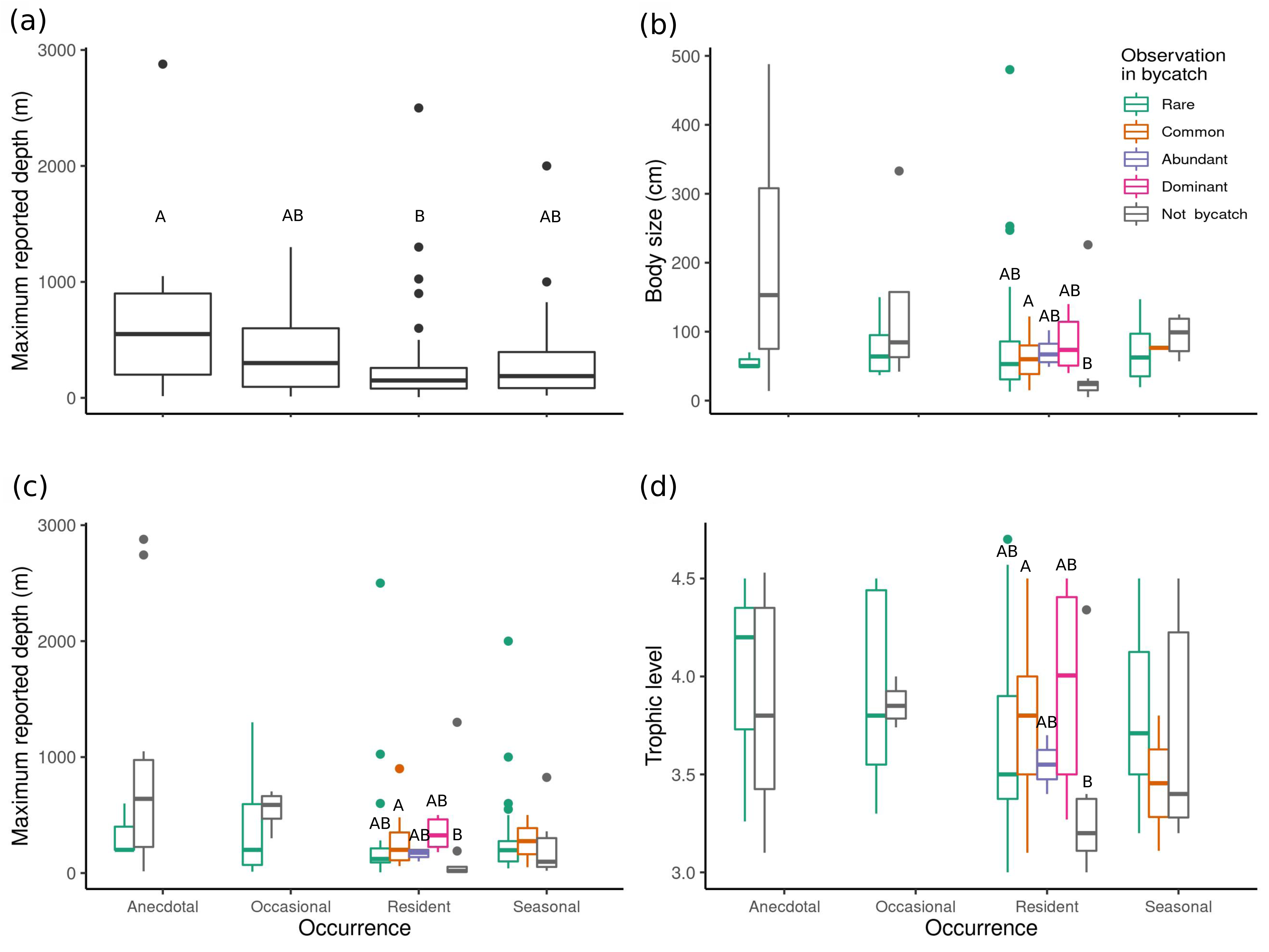
Significant differences in continuous trophic traits of fish species according to their occurrence status in the study area (a) and combination with observation percentage in the bycatch (b,c,d). Lower and upper box boundaries for maximum depth (a,c), body size (b), and trophic level (d) represent the 25th and 75th percentiles, respectively. The line inside the box is the median. Lower and upper error vertical lines represent the 10th and 90th percentiles, respectively, and coloured dots are data falling outside 10th and 90th percentiles. Uppercase letters above boxes denote significant differences in trait values of fish species among categories of bycatch inclusion per occurrence status.

### Spatial variation in fish functional diversity

Reference values of functional trait diversity calculated for the reconstructed fish assemblage and in the bycatch showed a low functional trait redundancy. The 127 fish species recorded represented 122 functional entities [roles], which left a redundancy of 1.04 fish species on average to accomplish a role in the entire area. Functional richness for this assemblage equals 1. The bycatch functional entities ranged from one to 28, with an average of eight, and functional redundancy ranged from 1 to 1.28, one fish species per functional role.

As we hypothesised, the most parsimonious GLM models showed that predictors of the fleet dynamics related to changes in the fish taxonomic richness and richness and dispersion of functional traits in the bycatch (Figures 4; see Table S5 (wAIC = 1, Variable importance = 1), and S6-8). Latitude is slightly positively related to fish species richness and negative with functional richness and dispersion (Figure 4a,e, i). Depth was associated slightly with increases in fish species and functional richness and decreases in functional traits’ dispersion (Figure 4b,f,j). Fishing intensity showed a slightly antagonistic relationship with the three fish diversity indices (Figure 4 c,g,k). Furthermore, the strong slopes of the temporal use of fishing grounds are related to increases in fish species and functional richness and functional trait homogenisation in the bycatch (Figure 4 d,h,l).

**Figure 4.**
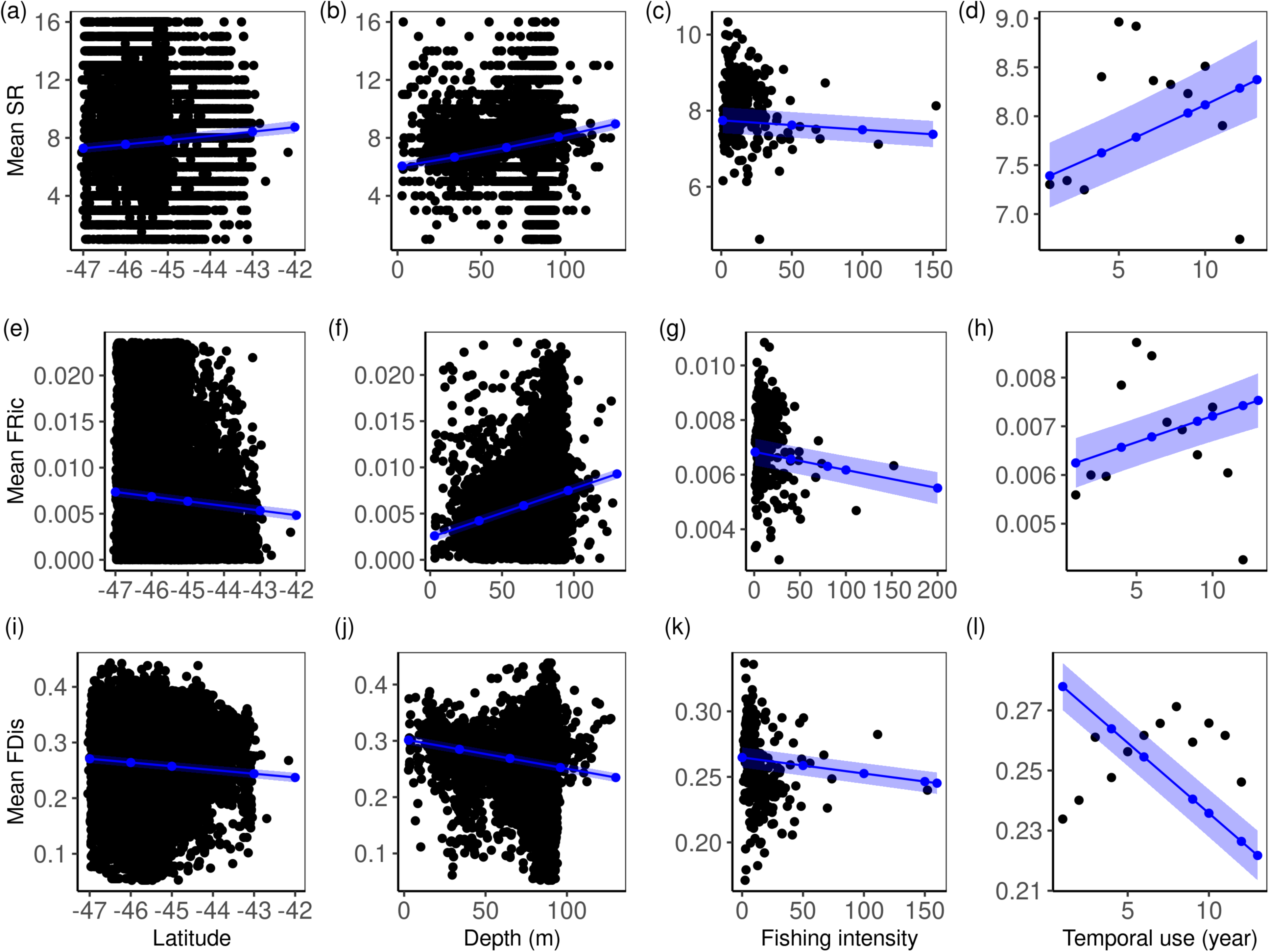
Relationship between fish taxonomic and functional diversity metrics in the bycatch with predictor variables. Relationships were plotted only for most parsimonious models of fish taxonomic richness (SR; a,b,c,d), and functional richness (FRic; e,f,g,h), and dispersion (FDis; g,h,i,j) with wAIC and variable importance equals to 1. Black dots denote observed mean values of diversity metrics. The solid blue lines and areas represented estimates and 95% confidence intervals from models, respectively.

### Spatial changes in fish trait community composition in the bycatch

The bycatch’s community mean trait values revealed differences in the dominance of water column positions and body shapes along gradients of predictor variables. Benthic-demersal, demersal, demersal-pelagic, and benthopelagic fishes had wider latitudinal (-44□ to -47□ S), bathymetric (10 to 120 m depth), and trawling intensity ranges (1 to 150 hauls/cell) than other positions in the water column (Figure 5a-c). Also, fishes that use benthic to pelagic areas dominated the bycatch at fishing grounds with higher temporal use (medians = 4 to 6 years) than pelagic-neritic fishes (median = 7.5 years, Figure 5d). By considering fish body shapes, we found that anguilliform, depressiform and elongated fishes dominated the bycatch at wider latitudinal ranges and deeper grounds (-44□ to -47□ S, and more than 75 m) than compressiform, fusiform, and fusiform-compressed fishes (-45□ and -46□ S, and below 80 m; Figure 5e,f). All body shapes dominated the bycatch below 50 hauls per cell, but their medians showed a pattern of dominance from low to intermediate fishing intensity by depressiform and elongated, anguilliform and compressiform, and fusiform and fusiform-compressed fishes (Figure 5h). By considering medians of temporal use, compressiform fishes dominated the bycatch in fished areas with less than three years of use, followed by elongated, anguilliform, depressiform, and fusiform, and fusiform-compressed (Figure 5h).

**Figure 5.**
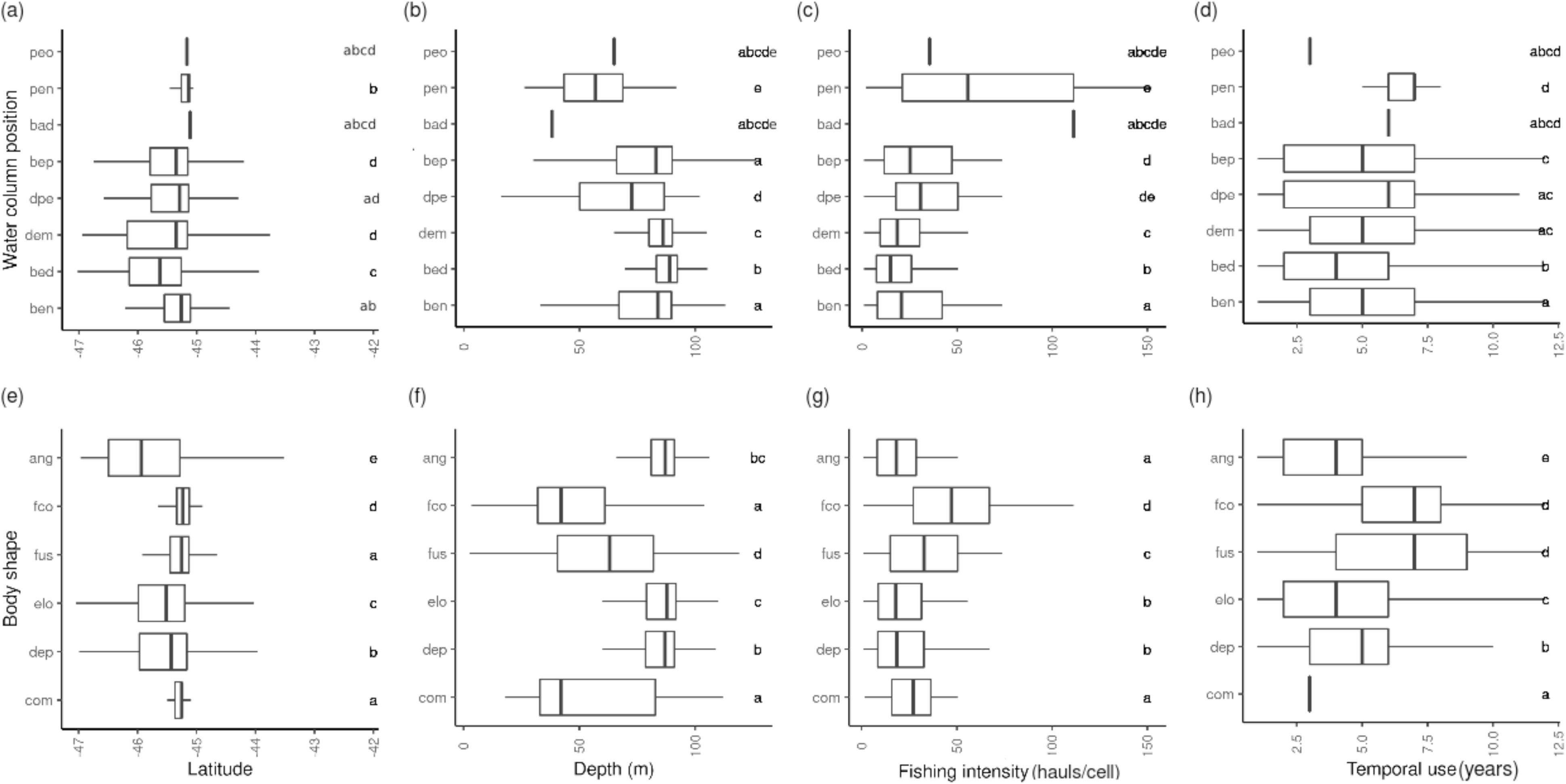
Spatial variation of dominant categorical fish functional traits in the double-rigged otter trawler fleet’s bycatch between 2003 and 2014. Letters denote significant differences in predictor values among trait categories (Mann-Whitney pairwise tests, corrected Bonferroni p-values < 0.05).

The most parsimonious models showed negative relationships between all fish community mean values of continuous traits in the bycatch with latitude and temporal use of fishing grounds and variable responses with depth and fishing intensity (Figure 6). Regression slopes showed that the mean fish body size strongly increased with depth, decreased northwards, and slightly decreased with fishing intensity and temporal use of fishing grounds (Figures 6a-d; Table S9). Reproductive load decreased northwards, with depth, fishing intensity, and temporal use of fishing grounds (Figures 6e-h; Table S10). The maximum depth of fish assemblages strongly increased with fishing grounds’ depth and slightly decreased polewards and fishing intensity (Figures 6i-k; Table S11). Trophic levels strongly increased with depth and slightly decreased northwards and with temporal use of grounds (Figures 6l-o; Table S12). Community mean diet breadth decreased with latitude and depth (Figures 6p-r; Table S13). In general, our results showed the importance of evaluated predictors of fleet dynamics in fish functional trait changes in the bycatch.

**Figure 6.**
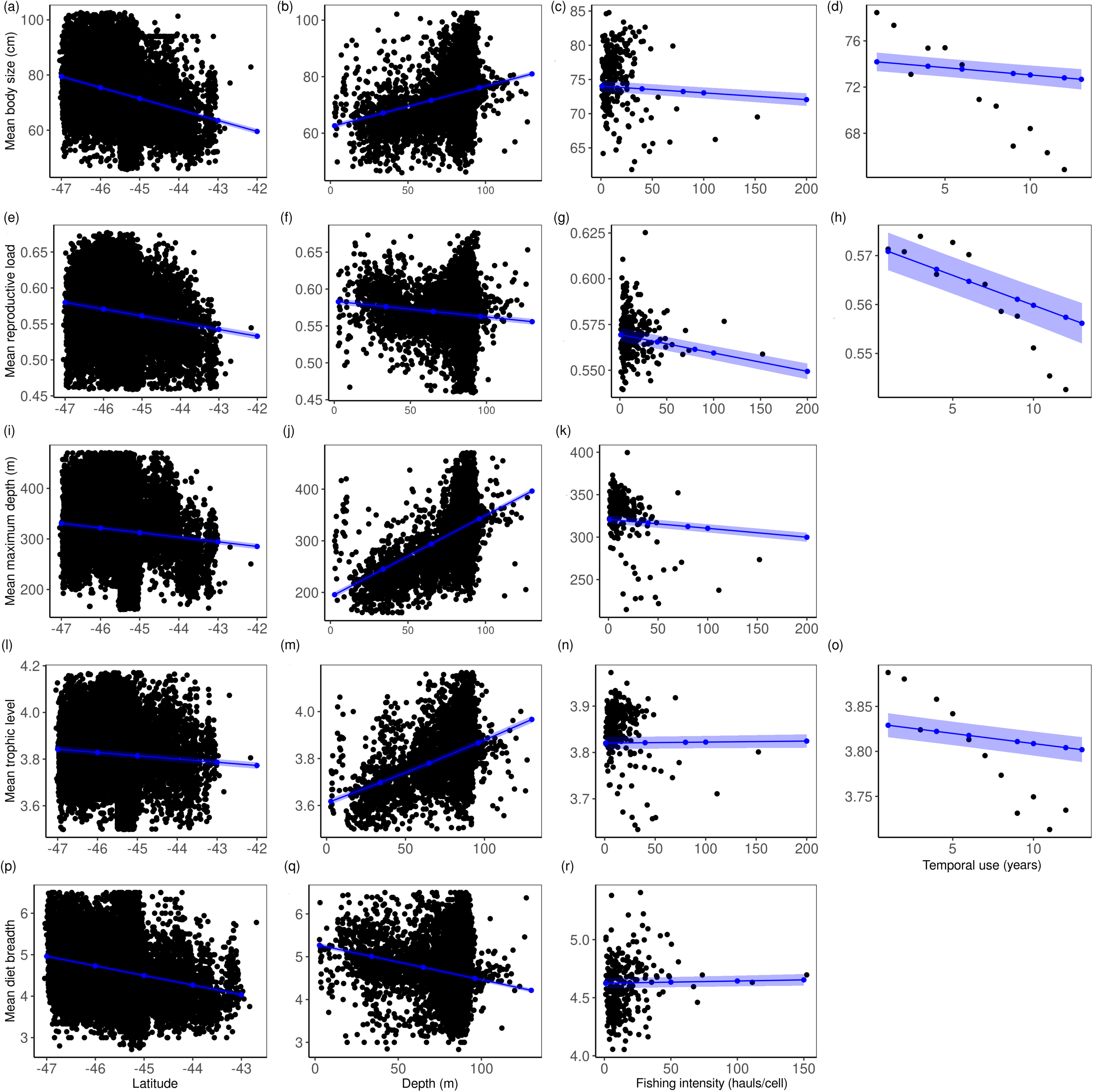
Relationship between fish community weighted mean trait values in the bycatch and predictor variables of the double-rigged otter trawler fleet between 2003 and 2014. The predictors of high importance for body size (a,b,c,d), reproductive load (e,f,g,h), maximum reported depth (i,j,k), trophic level (l,m,n,o), and diet breadth (p,q,r) were plotted (with wAIC and variable importance equals to 1). Black dots represent observed mean trait values, solid blue lines and areas estimates and 95% confidence intervals from models.

### Temporal variation in fish functional trait diversity metrics and composition

We observed significant declines in fish taxonomic and functional diversity and some community mean trait values after 2007 (Figure 7). The observed taxonomic and functional richness and diet breadth increased between 2003 and 2008 but then gradually declined with no return to higher values found in 2008. Mean fish community body size, reproductive load, and trophic level declined since 2003 with no return to baseline values. Fish body size declined 12 cm in length, reproductive load decreased 0.04 ratio, and trophic level 0.14 units on average along the 12 evaluated years. We also detected a 0.02 slight significant decrease in functional dispersion since 2003. No temporal change in the maximum depth of fish assemblages was observed. Evaluated random factors in the GLM models followed the decline in taxonomic and functional richness, but for the other diversity metrics and mean trait values, predictions remain stable despite their observed decline.

**Figure 7.**
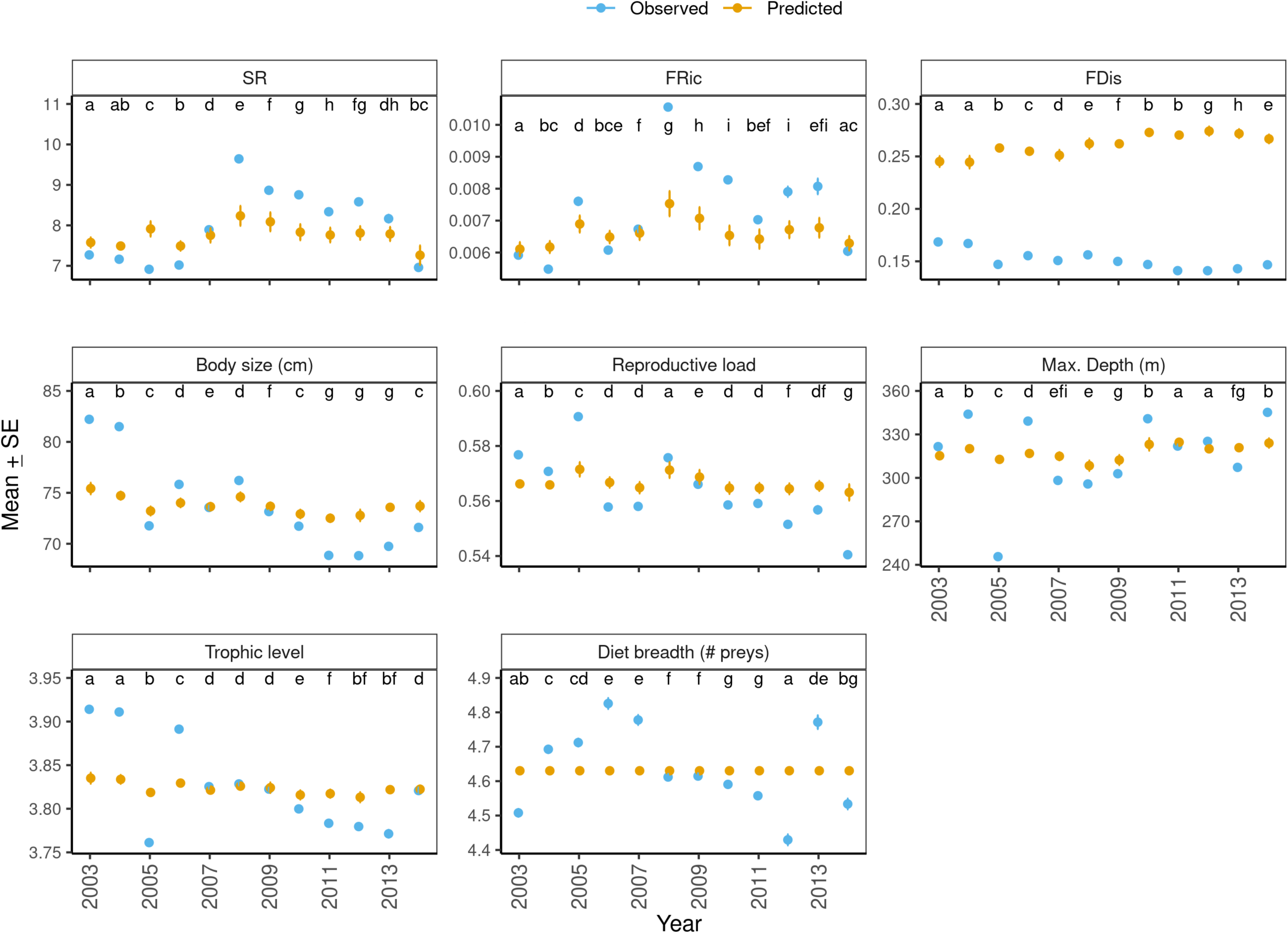
Temporal trends in fish species (SR) and functional richness (FRic), functional dispersion (FDis), and community weighted mean trait values of fish assemblages in the bycatch of the double-rigged otter trawler fleet. Means and standard errors (S.E.) were plotted for the annual mean observed values (blue dots) and model estimates (orange dots).

## Discussion

Evaluating fisheries’ effects on marine biodiversity through trait-based approaches is a challenge in areas where they are novel and an opportunity to find metrics useful for improving ecosystem-based fisheries management. For the first time in the Argentine Sea, we coupled trait information of 127 marine fish species with 12-years of data on the bycatch produced by the *P. muelleri* fishery and used three functional diversity metrics, and community mean trait values as descriptors of fish assemblage resilience to the industrial fishing. We found high functional trait diversity with low redundancy for the entire study area, in which each ecological role was accomplished on average by one fish species. Marine fish functional diversity, traditionally referred to as functional group richness in global models, decreases polewards, with two or three functional groups in coastal rocky reef systems in Argentina’s central Patagonia (Stuart-Smith et al., 2013). Using similar traits used in global models, we identified a higher richness of functional roles and considered four explanations.

First, our reconstructed fish assemblage included species from both coastal and marine areas up to 120 m depth within the trawler fleet’s operational area. This wide bathymetric range would allow other fish functional traits, increasing functional roles in the entire area. Second, the San Jorge Gulf (SJG) and adjacent waters are within the ecotone of two regional marine biogeographic zones in the Southwest Atlantic (-43° to -46° S), that receive species from subtropical and sub-Antarctic origins (Balech and Erlich, 2008), influencing also fish species composition in the bycatch (Bovcon 2016; Gongora et al., 2020; Figure 1a). Third, since the 1990s, the study area experienced the influx, range expansion, and establishment of 24 temperate-warm fishes from the Argentine Biogeographic Province, circumtropical and Brazilian areas that coincide with an increasing sea surface temperature north of -48° S, suggesting that a tropicalisation of fish assemblages would have started in these areas (Bovcon et al., 2011, 2016, Galván et al., 2005, *in press*; Góngora et al., 2009; Trobbiani et al., 2013; Venerus et al., 2006). Our anecdotal fish species showed deeper maximum depths than residents in the study area. This result aligns with the increasing fish functional trait richness found in unprotected temperate systems due to the tropicalisation of fish assemblages (Bates et al., 2014), which in our case implies a wide habitat distribution throughout the water column and deeper areas of the Argentinian shelf by newcomer species. We consider that the broad bathymetric range of our fish assemblage reconstruction, the influx of fish species from two distinct marine biogeographic areas and environments, and the initial tropicalisation of fish assemblages in this highly productive system within the Argentinian continental shelf increased fish functional trait diversity, but reduced their functional redundancy.

### Spatial drivers of fish functional diversity in the bycatch

Spatial fish trait variability does not necessarily couple patterns of species richness, and in contrast, may reflect changes in trait relative abundance or biomass in fish assemblages along gradients of human activities and environmental filtering (Dencker et al., 2017). Our results on fish species richness in the bycatch agreed with the expected increase of fish species northwards, but the richness and dispersion [variability] of functional roles decreased slightly at these latitudes, suggesting the inclusion of novel fish functional traits in the bycatch polewards. We observed a slight increase in mean fish body size, reproductive load, maximum reported depth, trophic level, and diet breadth polewards. While we still consider that species influx in this biogeographic ecotone may play an essential role in increasing fish species richness and functional roles polewards, the fleet’s spatiotemporal patterns and species abundance distribution in the bycatch would influence the observed spatial structure of fish functional diversity and trait composition.

The double-rigged otter trawler fleet’s spatial dynamics coincided with three seasonal thermal fronts located at the study area that may influence fish functional diversity. Observed grounds with high fishing intensity and temporal use at the northwest side of the SJG (around -45°S, - 66°W) and along the -66°W in its adjacent shelf (Figures 1b,c and S3), matched locations of local thermal fronts with high primary productivity and quality food for the shrimp *P. muelleri* (Figure 4 in Glembocki et al., 2015). Marine fronts in the South Western Atlantic are substantial fishing grounds and one of the leading spatial drivers in species compositions of local fish assemblages (Alemany et al., 2009, 2014). Highly productive areas support foraging of benthic and demersal invertebrates and promote benthopelagic coupling in food webs through food resources for demersal and pelagic fish species (Glembocki et al., 2015), forming functional niches for several species, including those in upper trophic levels such as sharks and rays. Location of thermal fronts coincided with areas of high dispersion [variability] in fish functional roles in the bycatch southern -45°S (Figure S6), explaining its latitudinal increase southwards compared with fishing grounds northwards.

The latitudinal structure of fish functional traits in the bycatch coincides with the spatiotemporal distribution of common cartilaginous species with intermediate to high trophic levels in the bycatch (>25% of observation), explaining our observed increased community body size, reproductive load, and trophic level polewards, and body size and trophic levels in deeper grounds (Figures S7-8, and S10). The otter trawler fleet incidentally caught chimaeras, sharks, and rays with more than 25% of observation in the bycatch of more than 5% of hauls deployed at northern grounds in the SJG and continuous deeper areas in national waters during the time of our evaluation (Ruibal Nuñez, 2020). Spatial captures coincide with records of the Plownose chimaera *Callorhinchus callorynchus* (Linnaeus, 1758); Narrownose smooth-hound *Mustelus schmitti* Springer 1939; Narrowmouthed catshark *Schroederichthys bivius* (Müller and Henle 1838); Picked dogfish *Squalus acanthias* Linnaeus 1758; Apron ray *Discopyge tschudii* Heckel 1846; Shortfin sand skate *Psammobatis normani* McEachran 1983; Smallthorn sand skate *Psammobatis rudis Günther 1870*; Smallnose fanskate *Sympterygia bonapartii Müller and Henle 1841;* Yellownose skate *Dipturus brevicaudatus* (Marini 1933); and Patagonian skate *Bathyraja macloviana* (Norman 1937) (see appendix A and B in Ruibal Nuñez, 2020). The low fecundity, considerable length at maturity in proportion to species maximum size, intermediate to high trophic level, benthic and demersal feeding habits, plus a high horizontal overlapping in the spatial distribution of these cartilaginous species with fishing grounds (> 30% of overlap) represented important traits that describe their vulnerability to this industrial fishing of low selectivity in our study area (Ruibal Nuñez, 2020). Besides supporting previous findings on the fish community’s vulnerable traits to this industrial fishing in our study area, our results also showed hot spots of functional trait erosion through the incidental capture of large predators.

Our results suggest that fishing grounds’ temporal use strongly correlated with fish functional diversity in the bycatch. We recognise that the temporal trends showed more complex behaviour than the linear, and 2008 may be a likely inflexion point in the functional diversity metrics and community trait values (Figures 4l and 7). Although another model type (e.g. GAM) might be more appropriate for treating these data, the GLMMs used here managed the variability of random factors, and initial results can be discussed and provide support for future working hypotheses on the system temporal dynamics. Overall, we observed a strong positive relationship between temporal use and fish species and functional richness and a negative slope with functional dispersion in the bycatch. These strong relationships showed that probabilities of removing different species and functional roles increase through time in a new fishing ground, but at a short-term use, trawling removed trait abundances reducing functional trait variability in fish assemblages. Indeed, we observed a decrease in fish mean community body size, reproductive load, and trophic level since a year of use of fishing grounds, and erosion of these traits in all the evaluated areas since 2003, the beginning of our evaluation. Global trends in fish functional diversity show that fisheries extract high levels of fish functional richness in many Large Marine Ecosystems, increasing particular functional groups and reducing functional evenness (Trindade-Santos et al., 2020). In our study, fish functional trait variability [dispersion] correlated positively with functional evenness and specialisation (Table S3). Since the mean fish functional trait variability decreased through time in our study area, we understand that assemblages would have experienced functional homogenisation by decreasing distinct functional trait combinations or specialist species with the cumulative use of trawled areas.

Low functional trait variability in marine fish assemblages captured by fisheries also relates to low levels of fish biomass productivity and the fishing industry’s resilience against climate variability (Dee et al., 2016). Swept area ratios of the otter trawler fleet calculated for northern areas of the SJG between 2013 and 2015 showed a high percentage of the area trawled (80%), but high fishing intensity areas represent only 2.3% of the fished area, with more than one haul per year in average (Trobbiani, 2018). Given that we determined the temporary use of a fishing ground with a minimum annual haul, it would be expected that the fleet’s intense and long-term swept areas, such as the gulf’s northern areas, show prolonged temporary erosions in functional trait dispersion. The observed low variability of fish functional roles in the bycatch suggests a low resilience capacity by fish assemblages to fished areas’ long-term use. This fact places temporal use as an essential predictor of spatiotemporal consequences of fishing in ecosystem functioning.

The fishing intensity was not a good predictor of fish functional diversity erosion as we observed slightly negative slopes of diversity metrics and community traits (body size, reproductive load, maximum depth) with the spatial fishing effort. Small increases of trophic level and diet breadth with fishing intensity were also identified. The other 85 fish species were abundant, common, and rare (n = 4, 21, and 60 species, respectively) with less than 50% observation (Gongora et al., 2020). The ecological role of rare species in the bycatch was unknown for our study area (Gongora 2011; Góngora et al., 2009, 2020). We identified functional trait ranges of rare fish species [body sizes, maximum reported depths, and trophic levels] wider than species with higher observation in the bycatch. This finding showed that rare fish species increase functional diversity through wide ranges of their trait values in the central Patagonian region. As the entire study area presents a low redundancy of functional roles, we think that the constant high number of rare species in the bycatch contributed to observe small changes in richness and variability of functional roles along the fishing intensity gradient.

We observed the depth of fishing grounds related to small increases in fish species and functional richness but reduced functional variability. Depth is a driver of fish community trait structure in temperate and subtropical northern regions and a proxy of environmental variables such as temperature, seasonality, and productivity (Dencker et al., 2017; Beukhof et al., 2019; Aguilar-Medrano and Vega-Cendejas 2019). In our study, deeper fishing grounds match highly productive thermal fronts at the southern coasts of the SJG and its deeper adjacent shelf, which could support increases in fish niche spaces in the bycatch. Fish community composition in the otter trawler fleet was dominated by six species, whose observation percentage increased with depth in our study area between 2003 and 2014 (*M. hubbsi* (97%), the Longtail southern cod *Patagonotothen ramsayi* (Regan 1913) (74%), Pink cusk-eel *Genypterus blacodes* (Forster 1801) (62%), *D. brevicaudatus* (55%), Southwest Atlantic butterfish *Stromateus brasiliensis* Fowler 1906 (54%), and Apron ray *Discopyge tschudii* Heckel 1846 (53%)) (Gongora et al., 2020). Ecosystem functioning expressed as fish biomass in temperate systems is not coupled with species richness but supported by the dominance of few generalist fish species with high trophic levels that use benthic and pelagic areas for feeding (Maureud et al., 2019a,b). This finding could explain the slight reduction in functional variability due to these species’ dominance in the trophic function. Besides latitude and temporal use of fishing grounds, depth worked as a good predictor of fish functional diversity in our area.

### Temporal changes in community trait composition in the bycatch

Functional diversity metrics and community weighted mean trait values are complementary tools to explain phase shifts in disturbed communities. We identify temporal erosion in the richness of species and functional roles after 2008, and their variability mainly by the 12 cm reduction in the fish community mean body size since 2003. This decrease seems to follow trends of fish assemblages in highly trawled areas in temperate regions. Long-term selective removal of fish functional traits such as large body size and high trophic levels led to trait homogenisation and novel trophic structures in fish assemblages due to bottom trawling in the North Sea (Barausse et al., 2011; Beukhof et al., 2019; Jennings et al., 2002). Our temporal reductions in fish community body size, reproductive ratio, and trophic level align well with the decreased community fish body size due to the high incidental capture of large cartilaginous fishes by trawling in the SJG in the last two decades (Funes, 2020; Ruibal Nuñez, 2020). Since the SJG is a hot spot for batoids in the Southwest Atlantic (Sabadin et al., 2020), our results recall the urge to protect the regional pool of Chondrichthyes (Gongora et al., 2020; Lucifora et al., 2012; Ruibal Nuñez, 2020), moreover because their functional extinction by high fishing pressure can trigger trophic cascades in ecosystems (Dulvy et al., 2014). Our findings suggest that changes in mean community body size and reproductive load were sensitive indicators of temporal changes in fish functional diversity caused by the industrial fishery of *P*. *muelleri*.

The observed decrease in the fish community mean trophic level contrasts with local increases previously found for two reef fish species and a complex trophic web in trawled areas of the SJG due to the inclusion of discards of high trophic level, such as *M. hubbsi,* in diets of intermediate predators (Funes, 2020; Funes et al., 2019). While we included fisheries discharge in the diet items, differences in trophic levels between studies may relate to differences in methods to calculate trophic levels and the number and taxon of marine species included. To better understand the effects of trawling on the functional diversity of fish assemblages in the area, evaluating trajectories of trait-based metrics between fished and non-fished areas with fisheries independent data is needed.

### Functional vulnerability to industrial fishing and further research needs

To achieve a balance between fishing activities and ecosystem functioning, the resilience of biotic communities potentially affected by these disturbances must be assessed. Functional trait-based approaches are assessment tools for fisheries and ecosystem-based management to clarify fish community resilience to multiple disturbances and ecosystem functioning changes (Barnett et al., 2019). Our work on spatiotemporal fish functional diversity changes is the first one on the Patagonian shelf and fills existing information gaps on this biodiversity component globally. Besides, we compiled information on seven functional traits synthesising almost 300 regional studies. This synthesis of information represents an important step to understand changes in ecosystem function and the resilience of fish communities to disturbances such as fishing or changes in oceanographic variables at the regional level of the South West Atlantic (SWA). Our study identified a low functional redundancy of fish assemblages in the SJG and adjacent shelf in a biogeographic ecotone with fish tropicalisation signs. Even though the trawler fleet removed fewer species and functional entities than our reference values for the entire study area, the bycatch removed unique trait combinations eroding their diversity. We showed that the cumulative temporal use of fishing grounds, with a minimum annual haul and year of use, is enough to erode the variability of fish functional traits exhibited by the bycatch’s body size, reproductive load, and trophic level.

The removal of functional traits from fish assemblages by trawling fleets worldwide appears to have no fixed selection pattern. However, we found essential indicators of fish species’ vulnerability to the double-rigged otter trawler fleet by identifying significant changes in trait categories between species in the bycatch and those recorded in the entire study area. By considering fish water column positions and body shapes, at least 50% of fishes in the bycatch use benthic, benthodemersal, and benthopelagic feeding areas and have depressed and fusiform shapes. We also identified vulnerable species to the fleet as resident fish species with body size above 30 cm TL, using habitats between 50 to 500 m depth and intermediate to higher trophic levels. Our vulnerable traits match the described taxonomic composition of dominant species and sharks and rays in the bycatch, whose observation percentage increases with depth (Gongora et al., 2020; Ruibal Nuñez 2020), and body shapes related to benthic/demersal feeding and living. In general, among evaluated traits, the water column position, body shape, body size, maximum reported depth, and trophic level provided important information on species’ vulnerability to trawling activities in our study area.

High vulnerability of large body-sized fish species to bottom trawling gears is documented worldwide (Beukhof et al., 2019; Henriques et al., 2014; Koutsidi et al., 2016; Pauly et al., 1998), as well as the increased capture of species with high trophic levels such as chondrichthyans by industrial fishing (Dulvy et al., 2017). We emphasise the importance of protecting rays and sharks in the study area because their functional characteristics match those described as vulnerable by this study. Their removal from the system and reduction in the bycatch are tipping points of observed changes in the local trophic structure. Even though the otter trawler fleet is the main one targeting the shrimp *P. muelleri*, at least two other fleets targeting the former one and *M. hubbsi* also incidentally capture chondrichthyans in the study area. While catches of multiple fisheries are skewed towards few (Mbaru et al., 2020) or multiple functional entities (Trinidade-Santos et al., 2020), the understanding of fish functional diversity of fisheries requires information from global captures instead of single fisheries (Nash et al., 2017). Further studies and local fishing managers may benefit from understanding all local fisheries’ combined effects in the spatial and temporal fish trait removal according to gear type to improve fisheries practices, management, and recovery of the area’s trophic function. Based on our evaluation, other information gaps need to be filled to improve the big picture of the fish functional diversity and resilience patterns at the regional level. First, a more comprehensive evaluation of fish functional diversity and its resilience must include both the bycatch and the system based on dependent and independent fisheries data. Observer programs onboard industrial fisheries have been vital for understanding regional fish biodiversity and its spatial and temporal changes in the SWA. The data provided by these programs need to be complemented with data from independent research cruises to assess the existing regional functional diversity and components removed by local fisheries. Second, our work characterised the abundance of functional traits by assigning an observation percentage within the bycatch. While this characterisation allowed us to glimpse changes in the spatiotemporal patterns of fish functional diversity consistent with previous changes in the trophic structure described for the area, it would be essential to collect information on the fish species biomass extracted by fisheries. Fish biomass changes can provide better information on ecosystem services such as food production and match data collected by fisheries independent cruises allowing comparisons of functional diversity metrics that use relative trait abundance such as dispersion, which changed in this study spatiotemporally. Third, our results seem to show a relationship between high functional diversity (richness and dispersion) with the limits of the study area’s local thermal fronts. Given that thermal fronts in the SWA coast in Argentina explain the marine fish biodiversity and productivity patterns (Lucifora et al., 2012, Alemany et al., 2014; Glembocki et al., 2015), including the fishing grounds proximity to these fronts as an explanatory variable is vital to clarify their influence in fish trait diversity and resilience to fishing activities. If thermal fronts and deeper areas support hot spots of fish functional trait richness and variability, these areas can improve ecosystem resilience and deserve special attention for protection and management.

## Acknowledgements

This work was funded by the fellowship for postdoctoral studies in strategic research topics of the Consejo Nacional de Investigaciones Científicas y Técnicas (CONICET), and Agencia Nacional de Promoción Científica y Tecnológica of the Argentine government (PICT 2017-2403). The Fishery Onboard Observer Program from the Fisheries Secretariat of the Chubut Province, Argentina, provided the bycatch data. Fish species in catches of the Argentine red shrimp fishery were registered by Alberto Pereyra, Carlos Moran, Cristian Cornejo, Diego Diaz, Gabriel Alonso, German Arenas, Gonzalo Quiroga, Guillermo Suarez, Jose Alvarez, Jose Castagno, Jose Fernandez, Juan Romero, Juan Linares, Juan Lastra, Juan Jimenez, Juan Torres, Julio Maier, Leonardo Jerez, Luis Villagran, Marcelo Olariaga, Marcelo Schmidt, María Vucica, Mariangeles Lopez, Matias Soutric, Mauricio Gallardo, Nelson Toledo, Nestor Santibañez, Osvaldo Muñoz, Pablo Evans, Rodrigo Cardenas, Rodrigo Torres, and Ruben Cambursano. We recognise Atila D.E. Gosztonyi, former curator of the ichthyologic collection at the CCT-Centro Nacional Patagónico (CENPAT) from the National Scientific and Technical Research Council of Argentina (CONICET), for his invaluable references used in the fish species reconstruction. Manuela Funes, Agustín De Wysiecki, Alejandro Irigoyen and Leonardo Venerus from the CONICET-Centro para el Estudio de Sistemas Marinos provided several references and personal observations used to compile the fish occurrences and functional traits.

## Contributions

MPRD and DEG conceived and designed the study. MPRD. conducted the literature review and data compilation of fish functional traits, verified the functional traits, conducted data analysis, and wrote this manuscript. NDB, MEG, and PDC provided fish bycatch data. NDB, PDC, and MPRD. compiled and verified fish species occurrences and their status. All co-authors provided editorial advice.

## Significance statement

Fish traits relate their functional diversity to vulnerability to industrial fishing. In the Argentine Patagonian sea, these relationships are scarcely studied to identify fish functional diversity spatiotemporal patterns and fishing levels at which these baselines shift. Since the first year of trawling for the shrimp *P. muelleri*, fish functional trait variability decreased in fishing grounds. This study highlights the temporal use of fished areas as an essential predictor of ecosystem functioning changes.

## Notes

Declaration of interest statement: The authors declare that there are no conflicts of interest.

### Competing Interest Statement

The authors have declared no competing interest.

